# A Polarized Histamine-GABA Core-Rim Architecture within Synaptic Vesicles

**DOI:** 10.64898/2026.08.05.743008

**Authors:** Kunio Fujiwara

**Author notes:** Mail address.

## Abstract

Neuroscience traditionally assumes that amino acid transmitters occupy clear synaptic vesicles, whereas monoamines reside in dense-core vesicles. Using a glutaraldehyde–NaBH₄ epitope-engineering platform enabling ultrastructural detection of small amines, we identify a polarized histamine–GABA vesicular organization within conventional GABAergic vesicles. Quantitative electron microscopy demonstrates histamine condensed into a dense intraluminal core, while complementary GABA immunolabeling supports the localization of GABA toward the vesicle periphery, consistent with a membrane-proximal rim. This conserved architecture across central, autonomic, and endocrine GABAergic systems provides a structural framework for temporally differentiated inhibitory signaling, challenges the clear-versus-dense-core vesicle paradigm, and establishes a unified principle for dual-transmitter architecture.

**One-sentence summary:** Using glutaraldehyde-NaBH4-based ultrastructural analysis, we identified a novel “core histamine–rim GABA” vesicular architecture within GABAergic neurons, fundamentally redefining traditional models of dual-transmitter co-packaging and release dynamics.

## Introduction

For more than half a century, synaptic vesicles have been divided into two canonical classes: clear vesicles containing amino acid transmitters that mediate rapid synaptic inhibition (*1*), and dense-core vesicles storing monoamines that support slower, modulatory signaling (*2*). Within this framework, histamine has been regarded as a monoaminergic neuromodulator largely restricted to the tuberomammillary nucleus (TMN) (*3*), and its distribution outside classical hypothalamic pathways has remained experimentally inaccessible. The failure to detect histamine in ultrastructural analyses has been due to technical limitations. The chemical properties of histamine have hindered the development of direct immunohistochemical (IHC) detection. Early histidine decarboxylase (HDC) antibodies, raised against fetal enzymes (*3*), lacked the sensitivity to detect low-level expression in adult tissue. Using a unique glutaraldehyde–NaBH₄ epitope-engineering platform, we overcome these long-standing barriers and directly visualize histamine at the electron microscopic level. This methodological advance reveals a previously unrecognized polarized “core histamine–rim GABA” architecture within conventional GABAergic synaptic vesicles.

Strikingly, the core–rim architecture was consistently observed across all examined GABAergic systems—from central circuits to peripheral structures such as adrenal chromaffin cells and sympathetic ganglia—indicating a conserved vesicular organization across diverse inhibitory populations within the tissues examined. Although these findings do not establish an organism-wide organizational principle, they substantially expand the experimentally validated domain of histamine/GABA co-packaging. Together, these observations challenge the long-standing segregation of amino acid and monoamine transmitters into distinct vesicle classes and establish a structural framework for histamine/GABA cotransmission.

## Results and Discussion

### Identification of a Polarized Core–Rim Vesicular Architecture

Electron microscopy identified a previously unrecognized polarized core–rim architecture within synaptic vesicles of conventional GABAergic neurons (Figs. 1 (*A*) and 2). This organization consists of a dense histamine core surrounded by the well-established peripheral GABA rim (*4,5*), thereby challenging the classical dichotomy between clear synaptic vesicles and monoaminergic dense-core vesicles. By segregating histamine and GABA into distinct intravesicular domains, the core–rim architecture represents a previously unrecognized mode of dual-transmitter organization. The histamine core was validated by exhaustive controls, including isotype controls, preabsorption, and primary antibody omission, all of which completely abolished diaminobenzidine (DAB) deposition. Because amino acid transmitters do not form dense cores under these fixation conditions (*1*), the polarized core–rim architecture represents a previously unrecognized structural principle for histamine/GABA co-packaging. This core–rim architecture extends the paradigm previously established in gastric enterochromaffin-like (ECL) paraneurons using the identical HA2 antibody (*6–9*). This conservation across CNS and endocrine systems aligns with the widely observed storage principle of other monoamines (*2*), which condense into dense cores for stable packaging. By positioning a histamine core within a GABAergic rim, this vesicular logic bridges the structural gap between monoaminergic sequestration and rapid synaptic transmission architectures (*1, 2*).

**Figure 1.**
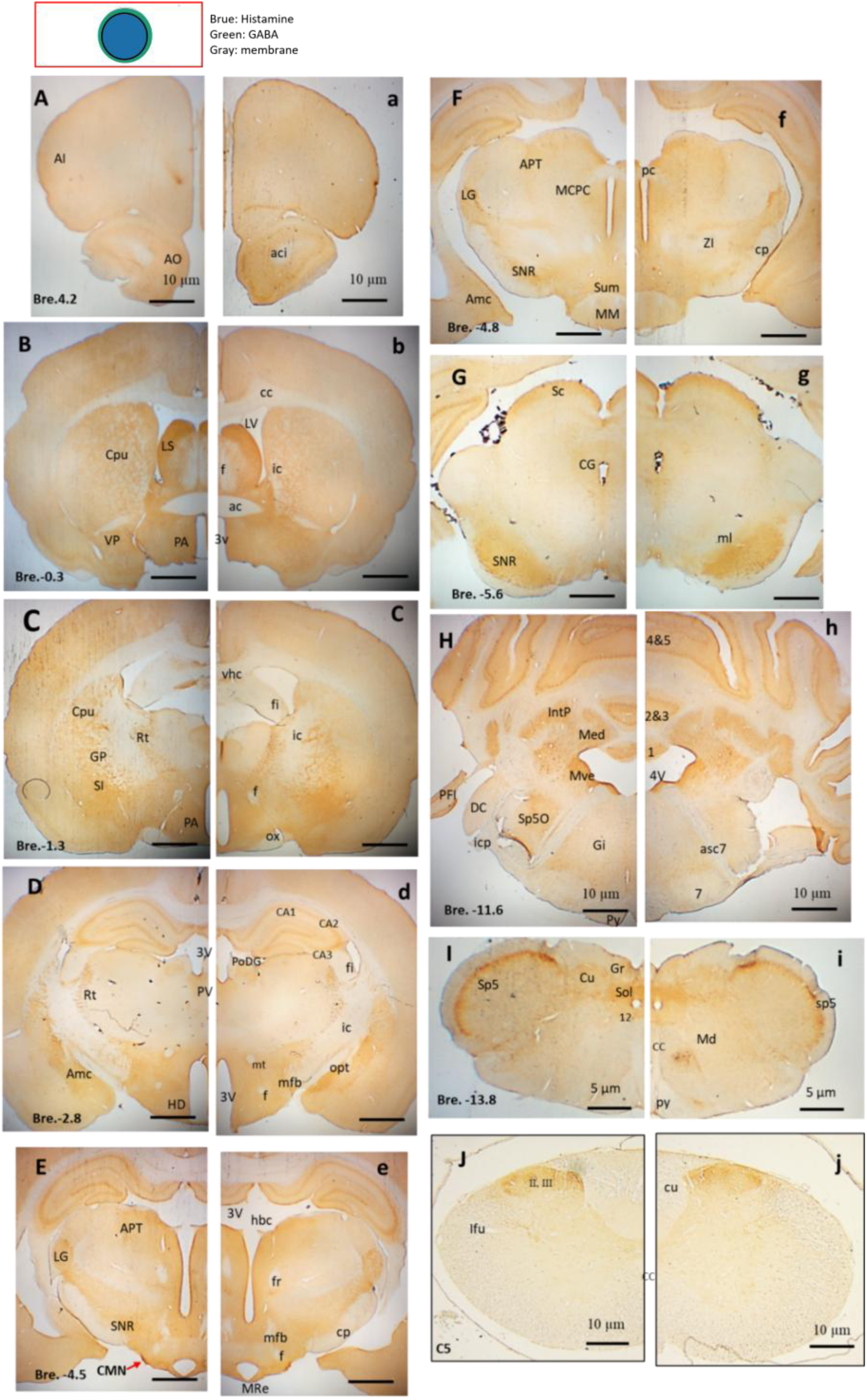
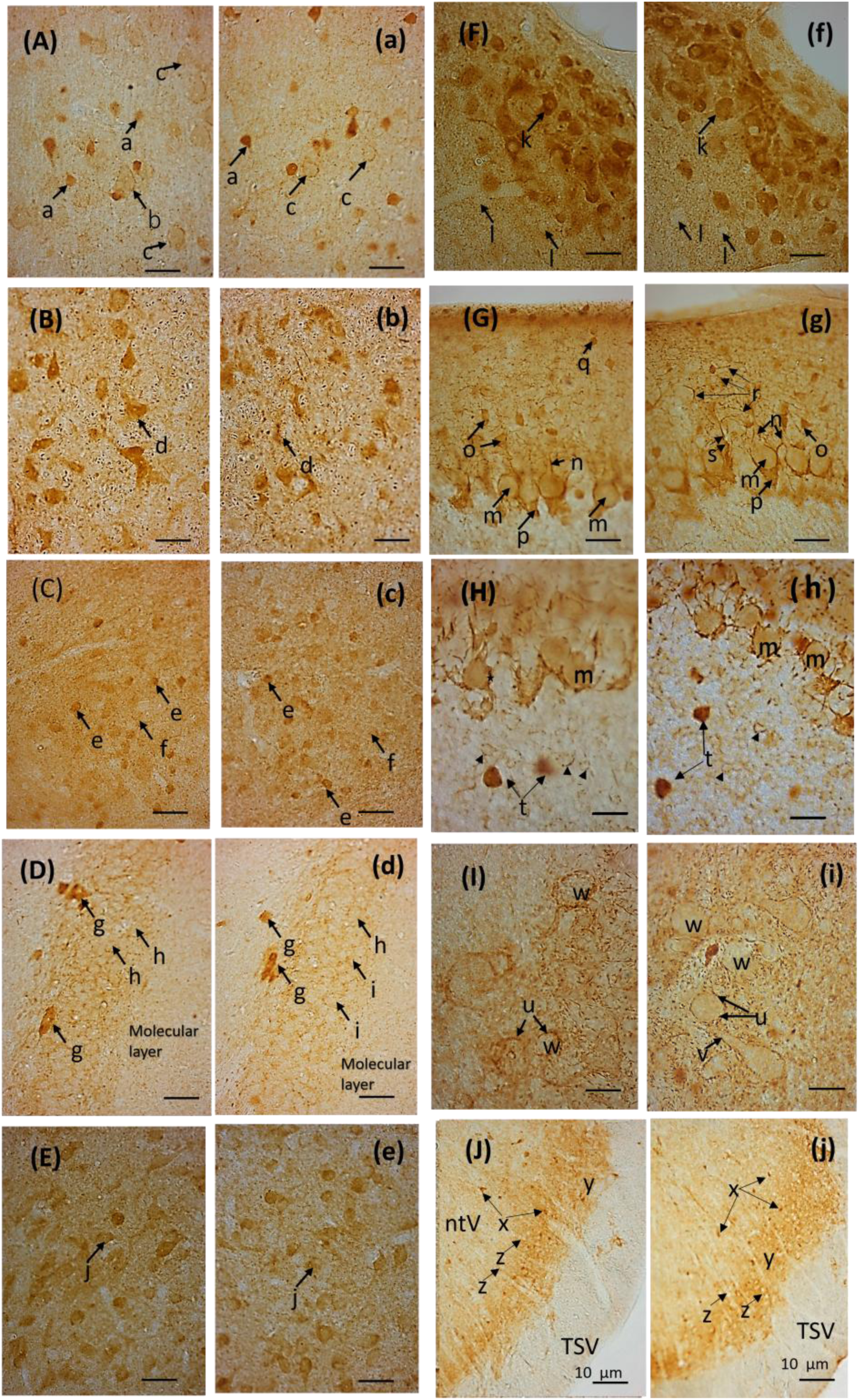
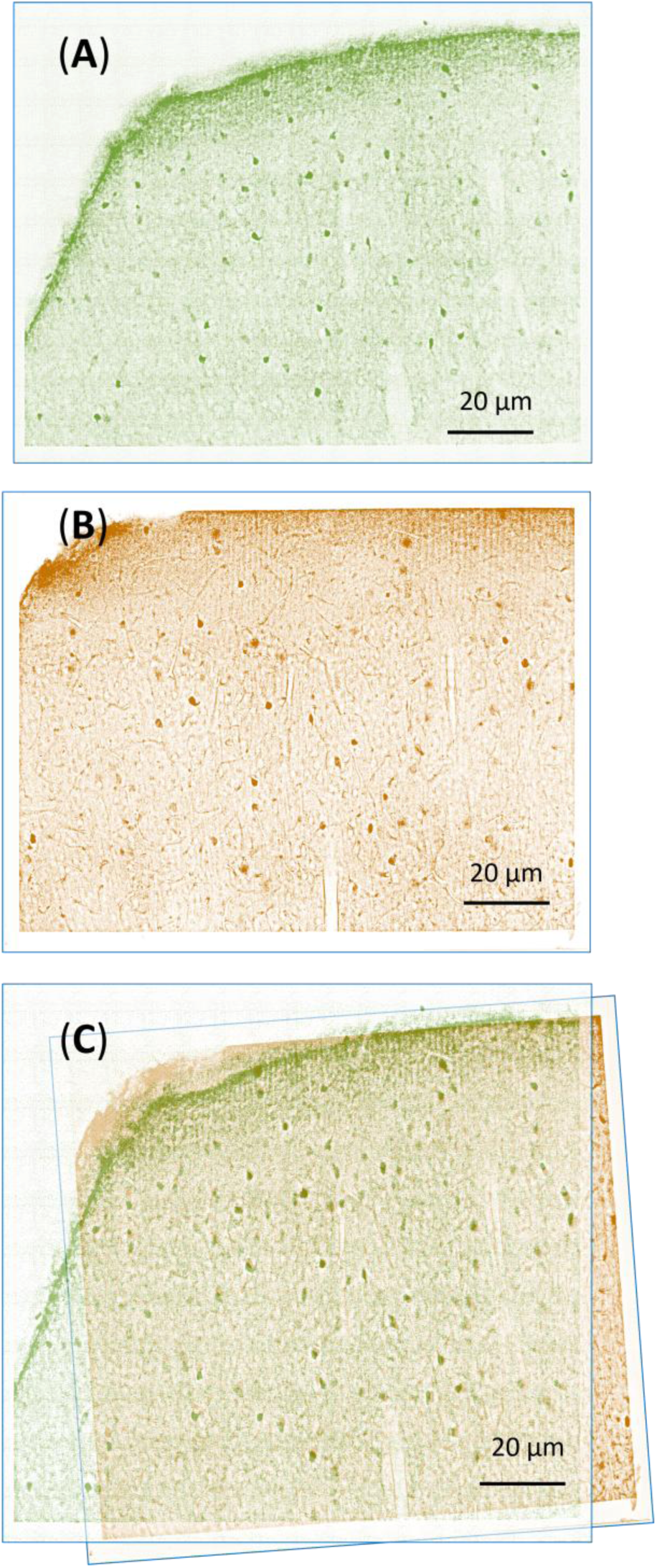
Core–rim organization and histamine/GABA colocalization in inhibitory neurons. This figure is organized into four hierarchical panels: (***A***) the core–rim structural model, (***B***) whole-brain distribution, (***C***) high-magnification cellular profiles, and (***D***) histamine–GABA colocalization. Together, these panels provide the light-microscopic foundation for the polarized core–rim architecture revealed by electron microscopy. **(A) Schematic representation of the core–rim vesicular organization.** A two-layer schematic illustrates the nanoscale segregation of transmitters within a single vesicle: the inner blue core marks the dense histamine-rich compartment, while the outer green rim denotes the membrane-proximal zone containing GABA. This diagram highlights their spatial separation. **(B) Distribution of histamine (A–J) and GABA (a–j) immunoreactivity in serial frontal sections processed under identical conditions.** Histamine-positive elements exhibit a spatial organization that closely parallels the GABAergic system (*30*), with anatomical localization referenced to the standard Paxinos and Watson stereotaxic atlas (*31*). Panels **A–J** and **a–j** are correspond to frontal sections at the indicated Bregma levels. **Abbreviations**: AI, agranular insular cortex; Amc, amygdaloid nu complex; AO, anterior olfactory nu; APT, ac, anterior commissure; anterior pretectal nu; aci, anterior commissure, intrabulbar part; asc7, ascending fibers of the facial nerve; CA1-3, fields CA1-3 of Ammon’s horn; CMN, caudate magnocellular nu; Cpu, caudate putamen; CG, central gray; Cu, cuneate nu; cc, corpus callosum; cp, cerebral peduncle; CC, central canal; cu, cuneate fasciculus; DC, dorsal cochlear nu, oral; f, fornix; fi, fimbria hippocampus; fr, fasciculus retroflexus; Gi, giantcellular reticular nu; GP, globus pallidus; Gr, gracile nu; HD, hypothalamic nu; hbc, habenular commissure; Ifu, lateral funiculus spinal cord; IntP, interposed cerebellar nu; ic, internal capsule; icp, inferior cerebellar peduncle; LG, lateral geniculate nu; LS, lateral septal nu; LV, lateral ventricle; MCPC, magnocellular nu of the posterior commissure; Med, medial cerebellar nu; Md, medullary reticular nu; MM, mammillary nu; Mre, mammillary recess; MVe, medial vestibular nu; mfb, medial forebrain bundle; ml, medial lemniscus; mt, mammillothalamic tract; opt, optic tract; ox, optic chiasm; PA, preoptic area; PoDG, polymorph layer of the dentate gyrus; PV, paraventricular nu; PFI, paraflocculus; pc, posterior commissure; py, pyramidal tract; Rt, reticular thal nu; Sc, superior colliculus; SI, substantia innominate; SNR, substantia nigra reticular; Sol, nu solitary tract; Sp5, spinal trigeminal nu; Sp5O, spinal trigeminal nu; Sum, supramammillary nu; sp5, spinal trigeminal tract; VP, ventral pallidum; 3V, 3rd ventricle; 4V, 4th ventricle; vhc, ventral hip commissure; ZI, zona incerta; 7, facial nu; 12, hypoglossal nu; 1,2,3,4,5, cerebellar lobules; II, III spinal cord layers. Scale bars: 10 μm (**A-H**, **J**; **a-h, j)**, 5 mµ (**I**; **i**) **(C) Representative histamine (A–J) and GABA (a–j) immunoreactivity at higher magnification**. **(A, a)** Cerebral Cortex: Layer III–IV. HA/GABA-positive non-pyramidal cells (a) are observed. Intense punctate terminals outline the perikarya of large immunonegative (IN) pyramidal cells (b) and round cells (c). **(B, b)** Ventral Pallidum: Large pleomorphic neurons (d) exhibit robust immunoreactivity. Note the presence of numerous IP axial cylinders within myelinated fibers. **(C, c)** Amygdala: Medium-sized amygdaloid neurons show moderate (e) to weak (f) immunoreactivity. **(D, d)** Hippocampus: Highly IP polymorphic cells (g) are prominent. IN granule cells (h) are densely delineated by IP punctate dots (i). **(E, e)** Lateral Septal Nucleus: Moderate staining in medium-sized neurons (j) and their associated neuropil. **(F, f)** Caudal Magnocellular Nucleus (CMN): These neurons display the most intense immunoreactivity (k) in the brain. IP dots outline neighboring IN neurons (l) within the neuropil. **(G, g)** Cerebellar Paraflocculus (Molecular Layer): Purkinje cell bodies (m) and dendrites (n) are weakly labeled, whereas basket cells (o) and stellate cells (q) are strongly immunopositive. Characteristic pinceau formations (p), varicose puncta (r), and basket cell terminals (s) are evident. **(H, h)** Cerebellar Granular Layer: Strong IP staining in Golgi cells (t) and punctate profiles (arrowheads) at the periphery of cerebellar glomeruli. Granule cell clusters remain IN. **(I, i)** Deep Cerebellar Nucleus: IP bouton-like axon terminals (likely Purkinje-derived) punctately surround the somata (u) and proximal dendrites (v) of large IN neurons (w). **(J, j)** Spinal Trigeminal Nucleus: Robust immunoreactivity in cells (x) and neuropil (y) of the substantia gelatinosa. Neighboring cells (z) and the spinal trigeminal tract (TSV) show minimal reactivity. Scale bars: 2.5 µm (**A–I; a–i**); 10 µm (**J, j**) **(D) Colocalization of histamine and GABA in inhibitory neurons**. Double immunostaining demonstrates complete somatic colocalization of histamine and GABA in neocortical inhibitory neurons, with corresponding dendritic and terminal labeling (*30*). Neocortex, Bregma -1.3 mm; Scale bars: 20 μm **(A–C)**

### Quantitative Radial Validation of the Vesicular Core

Quantitative radial analysis confirmed marked histamine enrichment within the inner 30–50% of the vesicular radius, with 95.1% of the total signal confined to the core (*n* = 2195 vs. 108 vesicles across 23 boutons; Fig. 2. G–K). The absence of intraluminal membranes indicates that this nanoscale segregation is maintained by intrinsic physical mechanisms—such as matrix binding or charge-dependent condensation—rather than diffusion barriers. This geometrical arrangement provides a structural framework for temporally staggered transmitter release during a single exocytic event. Upon fusion pore opening, membrane-proximal GABA is expected to diffuse rapidly to mediate fast synaptic transmission. In contrast, based on the physical chemistry of histamine–matrix interactions (*11*), the condensed core component dissociates more slowly, yielding a delayed, sustained modulatory output. This distinct single-vesicle architecture strongly suggests that inhibitory networks coordinate and control neurotransmission across multiple timescales, consistent with endocrine histamine biology where our previous studies demonstrated strict HA2 confinement to gastric ECL cell granule cores (*7–9*).

**Figure 2.**
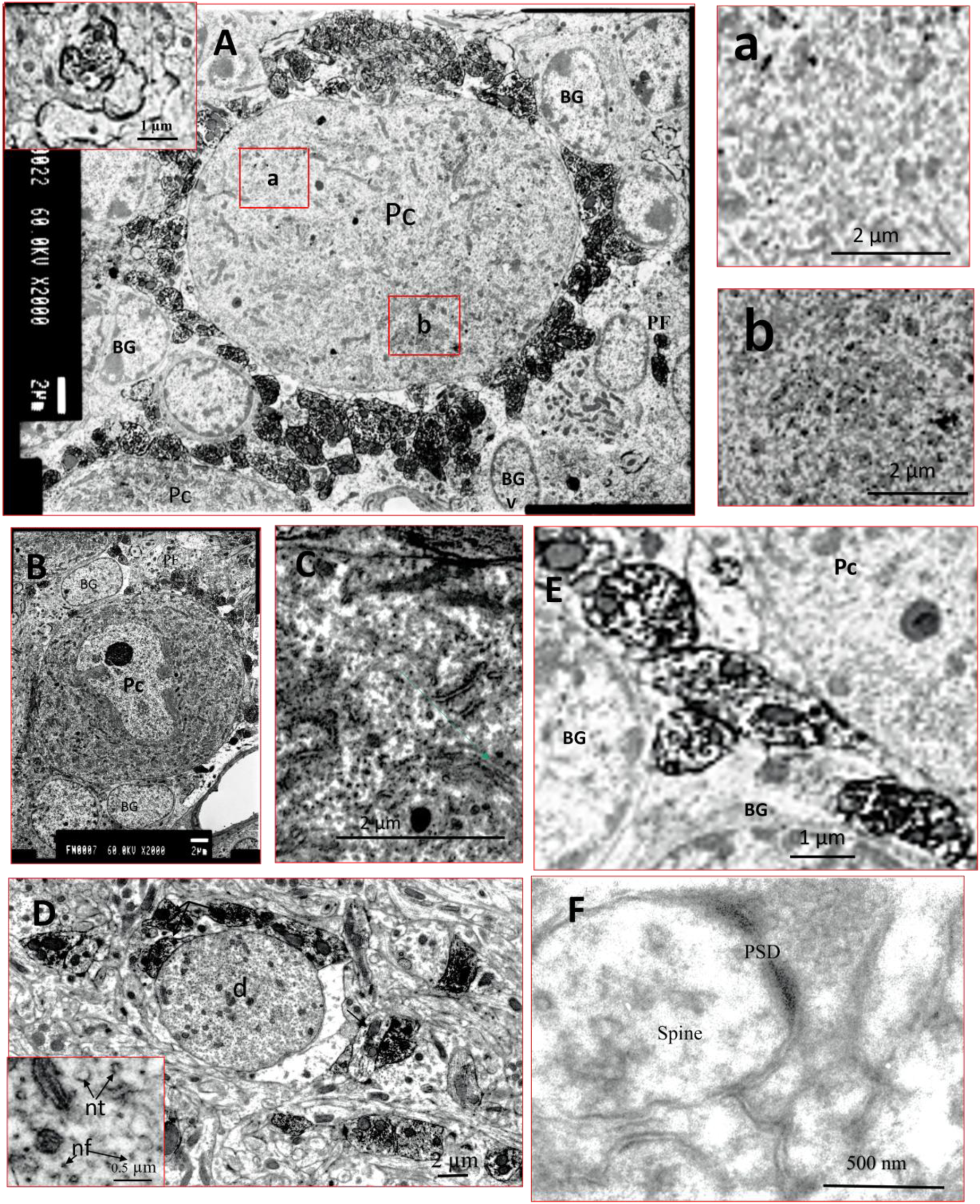
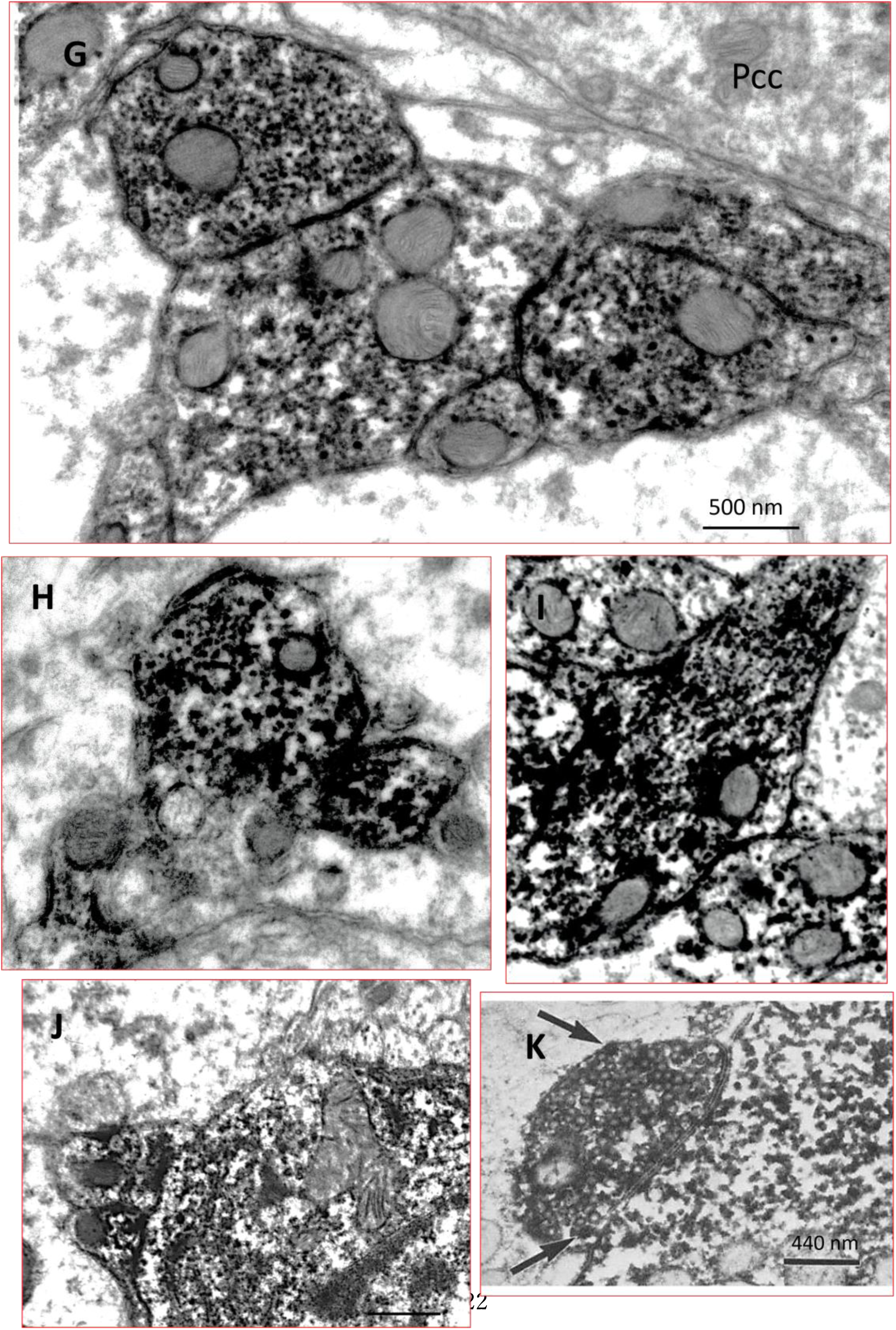
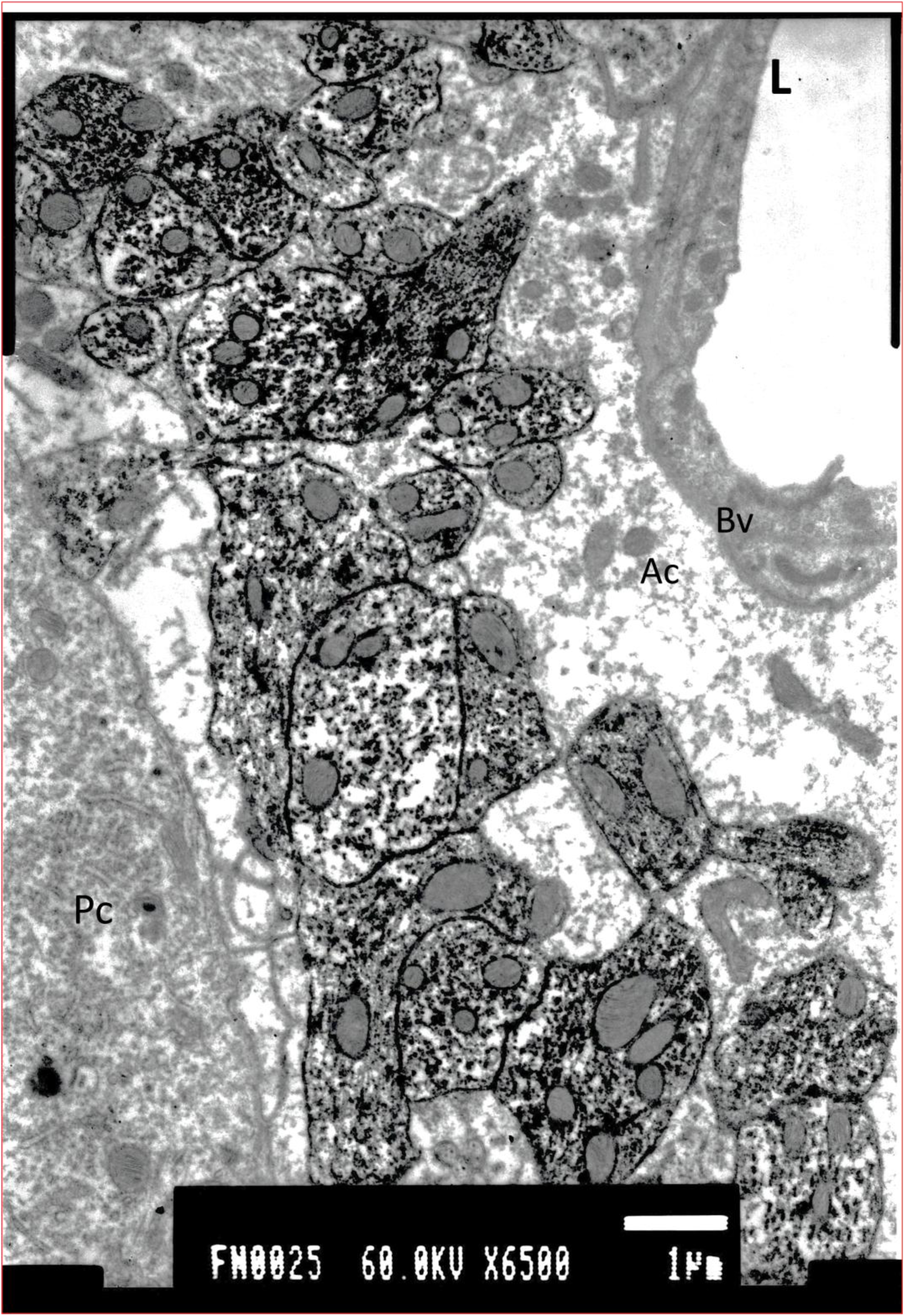
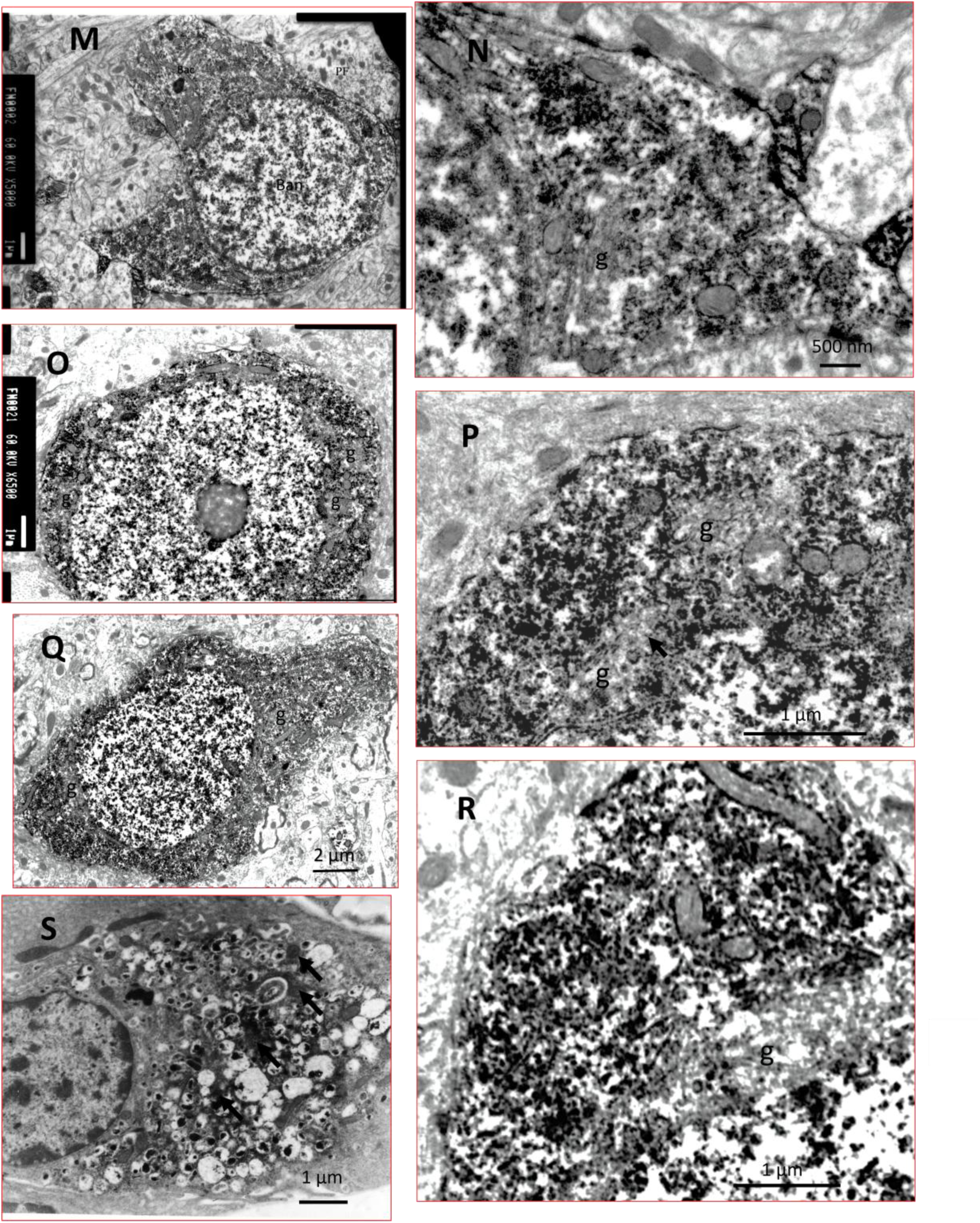
Electron microscopy of the polarized core–rim vesicular architecture. Four components: (**A–F**) low-magnification Purkinje-related ultrastructure, (**G–K**) high-magnification core–rim vesicles, (**L–R**) cell-type–specific inhibitory profiles, and (**S**) an endocrine reference—together establishing the ultrastructural basis of the histamine core and GABA rim. Electron microscopy revealed a polarized vesicular architecture in which histamine forms a dense central core, whereas GABA is confined to a thin peripheral rim. Apart from this core–rim organization, vesicles and synaptic morphology were indistinguishable from classical GABAergic terminals (*32*). Radial analysis demonstrated histamine enrichment within the inner 50% of the vesicle radius, and DAB quantification confirmed predominant core localization (95.1% of total signal; 2195 vs. 108 vesicles in 23 boutons; counts within Golgi regions excluded). **Purkinje cell layer (A–F).** Purkinje cell bodies and dendrites showed weak labeling, whereas ribosomes and short rough endoplasmic reticulum segments were strongly immunoreactive. Nerve filaments and microtubules were weakly labeled. Gray type I synapses containing round vesicles were immunonegative (IN), serving as internal controls (*33*). **High-magnification electron micrographs (G–K).** Basket cell terminals contained dense-core histamine-positive vesicles surrounded by a thin GABA-immunolabeled rim. Vesicles frequently aligned along microtubules (**I**). Occasional histamine-positive boutons formed asymmetric contacts with IN profiles (**J**). Panel (**K)** shows the classical peripheral GABA rim for comparison. **Medium magnification near Purkinje somata (L).** Basket cell collaterals formed symmetric Gray type II synapses with neighboring boutons but not with Purkinje somata. Vesicles consistently displayed central histamine labeling. **Basket, stellate, and Lugaro cells (M–R).** These neurons showed strong cytoplasmic labeling with clusters of histamine-positive vesicles around immunoreactive regions. Vesicles within the Golgi apparatus and mitochondrial matrix were unlabeled. **Comparative control (S).** Gastric ECL cells exhibited both dense-core and cytoplasmic HA2 labeling, providing an independent morphological reference. **Abbreviations:** Pc, Purkinje cell; Pcd, Purkinje dendrite; BG, Bergmann glia; Ac, astrocyte; g, Golgi region; d, dendrite. Differences in optical density between the vesicle core and peripheral rim were statistically significant (P < 0.001, Wilcoxon rank-sum test). Scale bars: 200 nm **(A–F),** 500 nm **(G–I, N),** 100 nm **(J),** 440 nm **(K),** 1 µm **(L, M, O, P, R, S),** 2 µm **(Q).**

### Technical Barriers Historically Obscuring Histamine Distribution

The observable range of neuronal histamine was previously limited to pathways originating from the tuberomamillary nucleus (TMN). The GA–NaBH₄ method expands this domain rather than revising established circuitry. In conventional models, neuronal histamine synthesis was considered largely restricted to TMN (*3,12*). This view originated from methodological limitations: early indirect immunohistochemistry for histidine decarboxylase (HDC) relied on antibodies raised against fetal enzymes (*3*), which may have lacked the sensitivity to detect low-level expression outside classical pathways, including atypical synthetic enzymes or previously unrecognized HDC isoforms. Likewise, early histamine mapping failed because weak formaldehyde fixation caused poor tissue retention (*12*).

### Technological breakthrough using the GA–NaBH₄ method

Antigen–antibody interactions generally require an epitope at least the size of a dipeptide or trisaccharide for stable molecular recognition (*13,14*). We developed an epitope-engineering strategy targeting aliphatic primary amines, a pioneering platform now extended for the first time to the ultrastructural localization of polyamines (*15*) and exogenous drugs (*16,17*). Because histamine is too small for direct antibody recognition, specificity depends on detecting the histamine–GA moiety generated within fixed protein complexes, the same chemistry underlying in situ fixation. Subsequent NaBH₄ reduction converts reactive aldehydes into non-reactive alcohols, markedly reducing non-specific background staining. The histamine–GA–BSA immunogen generates at least two GA-dependent epitope classes recognized by distinct monoclonal antibodies: HA1/3–5 and HA2. Whereas gastric ECL cells express both epitope classes (*6–9*), neural tissue appears to generate only the HA2-type epitope (*18*).

Retaining volatile histamine requires strong GA fixation, which severely restricts tissue permeability. Our DAB pre-embedding method overcomes this limitation by allowing antibodies to penetrate the vesicular lumen—where the immunoreaction and subsequent DAB deposition occur—thereby enabling high-resolution localization analysis at the ultrastructural level. In contrast, immunogold labeling is unsuitable for dual immunolabeling of histamine and GABA because colloidal gold particles cannot penetrate the vesicular membrane (*19, 20*).

**In the cerebral cortex,** at the light microscopic level, this permeability barrier was bypassed enabling high-resolution DAB double staining revealing near-complete spatial overlap between histamine- and GABA-producing cells. Quantification confirmed a 100% overlap (110 of 110 cells), despite minor tissue shrinkage typical of the free-floating method (Fig. 1 (*D*)).

The high technical precision of the GA–NaBH4 method offers a distinct advantage in re-evaluating vesicular storage within cell bodies. Specifically, by maximizing the signal-to-noise ratio, this method has revealed the phenomenon of “somatic vesicular storage”—a feature previously overlooked even in conventional GABAergic cells. It provides the first direct ultrastructural evidence demonstrating the localization of histamine-immunopositive (IP) synaptic vesicles within the cell bodies of atypical cell types, such as cerebellar Purkinje cells (Fig. 2B, C), basket cells (Fig. 2M, N), stellate cells (Fig. 2O, P), and Lugaro cells (Fig. 2Q, R), not to mention within their axons (Fig. 2I, L) and terminals (Fig. 2A, D, E, G, H, J, L).

### Spatial Co-Localization of GABA and Histamine

To evaluate whether these robust genetic profiles translate into the localized end-product histamine, we examined the macro-scale spatial distributions of both GABA and histamine using high-resolution light microscopy. Immunohistochemical evaluation of adult rat cerebellar sections revealed a distinct geometric profile shared between both molecular systems (Figs. 1 (*B*)). Within the cerebellar cortex, the interior of the Purkinje cell (PC) somata exhibited faint but detectable immunoreactivity for both GABA and histamine (Fig. 1 (*B*); G, H, g, h). In the deep cerebellar nuclei (DCN), immunoreactive terminals for both molecules outlined the outer periphery of large DCN projection neurons, forming a thin, clear pericellular outline (Fig. 1 (*B*); I, i). This spatial configuration—characterized by a faintly stained somatic interior and a moderately stained peripheral rim (arrows w)—was consistently replicated for both GABA and histamine, confirming the selective distribution of these dual transmitters into efferent terminal fields.

This central signature further aligns with peripheral, neural-crest–derived networks. In the adrenal medulla, histamine immunoreactivity was detected in chromaffin cells organized in cord-like clusters (Fig. 3a, b), reminiscent of well-documented peripheral GABA–adrenaline systems (*21*) and consistent with control sections (Fig. 3a’, b’). Similarly, in the sympathetic ganglia, histamine expression was localized within neuronal cell bodies and along arriving nerve fibers that form peripheral pericellular networks (Fig. 3c, d), mirroring the localized GABAergic morphology in peripheral ganglia (*22, 23*).

**Figure 3.**
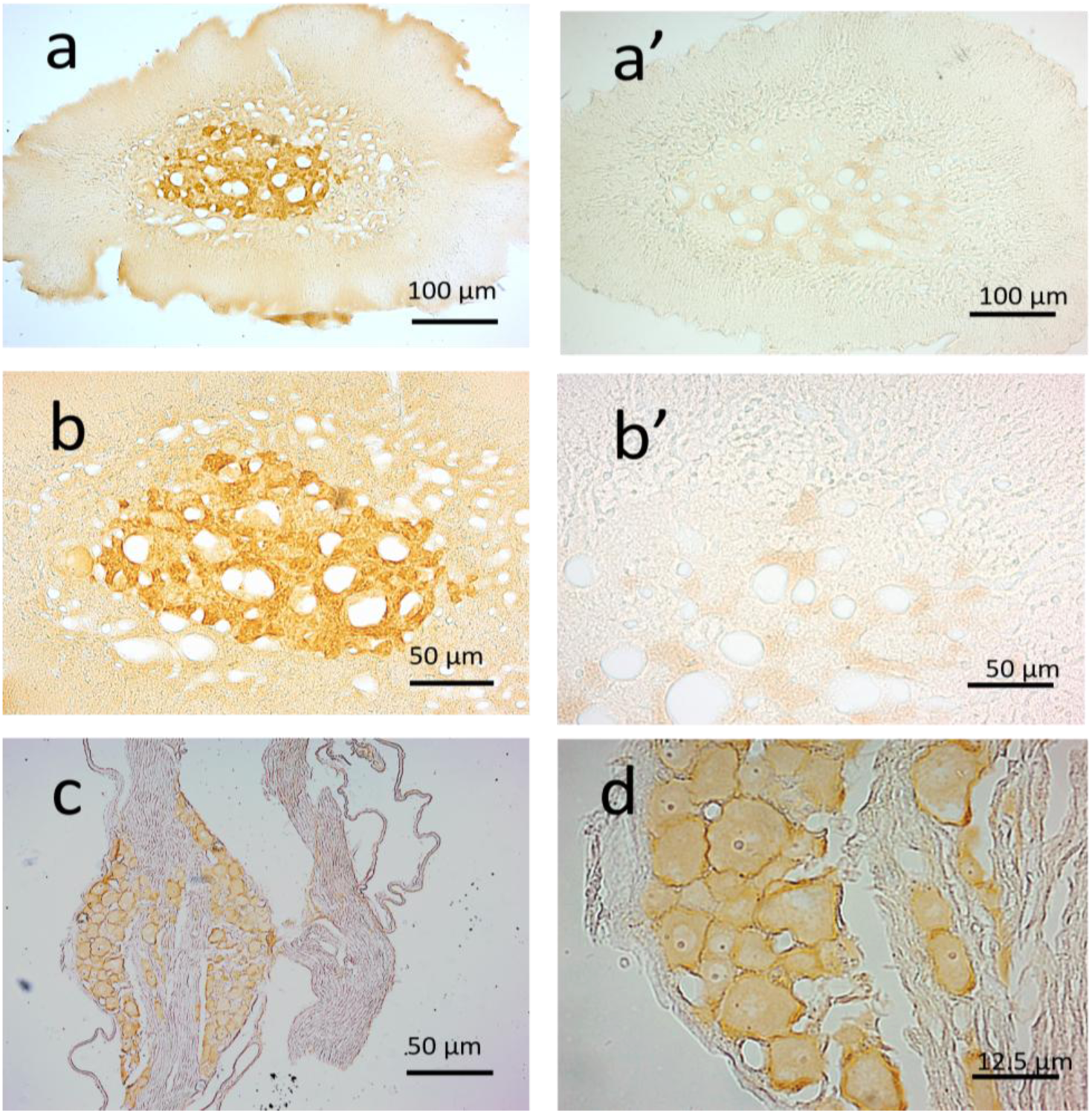
Conservation of the core–rim histamine/GABA organization in endocrine and autonomic systems. Three components: **(a–b)** chromaffin cell histamine storage, **(a′–b′)** HA2 absorption controls, and **(c–d)** sympathetic ganglia varicose fibers—showing conserved histamine/GABA core–rim organization in peripheral systems. **(a, b) Adrenal medulla.** Histamine immunoreactivity exhibits a punctate cytoplasmic pattern in chromaffin cells, consistent with ultrastructural evidence that histamine is stored in VMAT2-dependent dense-core vesicles and with classical studies of monoamine storage in endocrine cells (*21*). **(a′, b′) Absorption controls.** Preabsorption with histamine–GA–BSA and omission of the primary antibody both eliminated specific immunostaining. The faint residual coloration occasionally observed in noradrenaline (NA) cells was unaffected by either control, indicating that it reflects intrinsic oxidation products rather than incomplete antibody absorption or antibody-dependent staining. **(c, d) Sympathetic ganglia.** HA2-positive varicose terminals formed a dense pericellular network around principal neurons, whereas neuronal somata showed only weak immunoreactivity. Numerous histamine-positive fibers coursed between ganglionic cell bodies, forming a characteristic pericellular innervation pattern consistent with autonomic cotransmission (*22,23*). Together, these findings demonstrate that histamine/GABA cotransmission extends beyond the central nervous system to autonomic and endocrine tissues. Histamine-positive varicose fibers in sympathetic ganglia and VMAT2-dependent dense-core vesicles in adrenal chromaffin cells indicate a conserved vesicular organization across peripheral effector systems. **Scale bars:** 100 μm (**a, a′, c**), 50 μm (**b, b′**), 12.5 μm (**d**).

### *Hdc* Expression Analysis in Cerebellar Subclasses

Analysis of the Macosko cerebellar dataset revealed cell-type-specific expression of *Hdc* among the four major cerebellar subclasses (*24*). Purkinje cells exhibited the highest level of *Hdc* expression, with a mean expression of 0.912 and an *Hdc*-positive rate of 15.51% (130/838 cells). In contrast, molecular layer interneurons (MLIs) expressing *Megf11* (multiple EGF-like domains protein 11) showed moderate expression (mean: 0.149, 1.92% positive), whereas MLIs expressing *Cdh22* (cadherin-22) and Granule cells showed minimal to negligible expression (0.22% and 0.008% positive, respectively) (Fig. 4).

**Figure 4.**
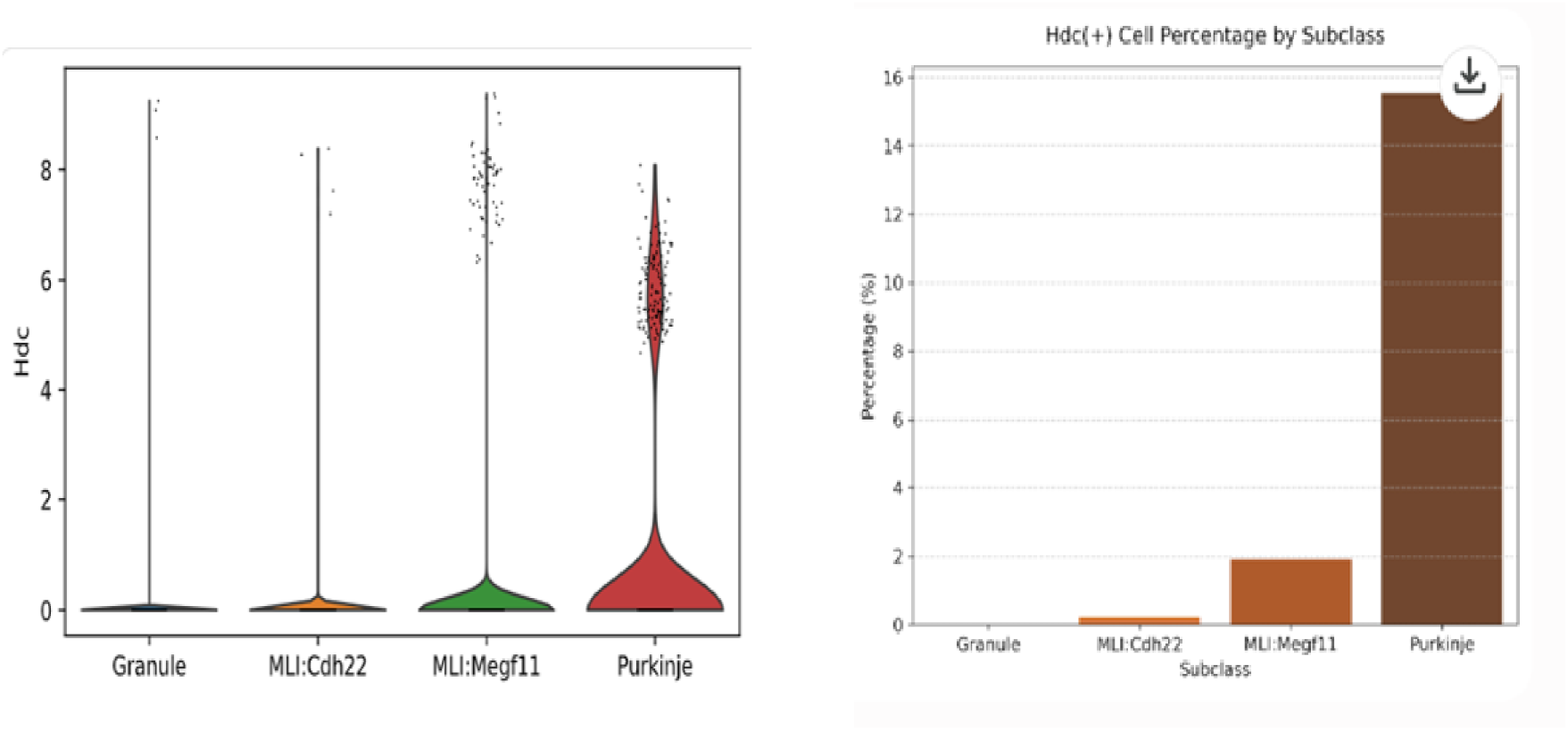
Analysis of *Hdc* expression and cellular proportion across cerebellar subclasses. Two components are shown: (Left) single-cell *Hdc* expression across cerebellar subclasses, and (Right) the proportion of *Hdc*-positive cells. Together, these panels indicate that Purkinje cells contain a distinct subset of *Hdc*-expressing neurons. **(Left)** Violin plot showing the distribution of *Hdc* expression levels in individual cells across four cerebellar subclasses: Granule, Molecular level Interneurons (MLIs expressing cadherin-22 (Cdh22), MLI expressing multiple EGF-like domains protein 11 (Megf11), and Purkinje. Each dot represents a single cell. **(Right)** Bar chart illustrating the percentage of *Hdc*-positive (*Hdc* (+)) cells within each subclass. The Purkinje cell subclass exhibits the highest proportion of *Hdc* (+) cells compared to the other groups.

### Functional Implications of Cell-Type-Specific *Hdc* Expression

The Allen (Macosko) dataset reveals low-frequency but unequivocal *Hdc* expression in Purkinje cells (15.5%), indicating that at least a subset of non-TMN neurons possesses the molecular machinery for histamine synthesis. The present study detects histamine, the final biosynthetic product, whereas the Allen dataset measures *Hdc* transcript abundance. Because these represent distinct biological layers, a quantitative correspondence is not expected. As in other monoaminergic systems, vesicular accumulation and recycling can maintain histamine levels independently of instantaneous *Hdc* mRNA abundance (*25,26*).

Furthermore, HDC is a PLP-dependent enzyme whose activity is influenced by cofactor availability, local metabolism, translational efficiency, and protein stability, further uncoupling transcript abundance from tissue histamine levels (*27,28*). The longstanding TMN-restricted model may also reflect the use of fetal-derived HDC antibodies in adult tissue (*3*), which may have underestimated low-frequency expression outside the TMN. Accordingly, the non-TMN histamine identified here does not contradict previous studies but instead expands the detectable histaminergic domain through the GA–NaBH₄ method. Together, these findings support a conserved histamine–GABA core–rim vesicular architecture underlying dual-transmitter organization Together, these findings support a conserved histamine–GABA core–rim vesicular architecture that extends the current framework of inhibitory neurotransmission.

### Conclusion: A Structural and Genetic Framework for the Core–Rim Architecture

In summary, the integration of an optimized immunohistochemical imaging platform with single-cell transcriptomics establishes a structural and genetic framework for the histamine–GABA core–rim architecture across inhibitory neural circuits. By overcoming longstanding technical limitations, our approach reveals histaminergic features in mature GABAergic neurons beyond the tuberomammillary nucleus (TMN), challenging the conventional TMN-restricted model of the histaminergic system.

Electron microscopy further identifies a polarized histamine–GABA core–rim architecture within synaptic vesicles, extending the classical dichotomy between clear synaptic vesicles and monoaminergic dense-core vesicles. Together, these findings establish the core–rim architecture as a previously unrecognized organizational principle of inhibitory neurotransmission and provide a unified structural framework for polarized histamine–GABA vesicular organization across inhibitory neural circuits.

Our approach reveals histamine localization in mature GABAergic neurons beyond the TMN.

## Figure Legends

**Extended Data Figure 1.**
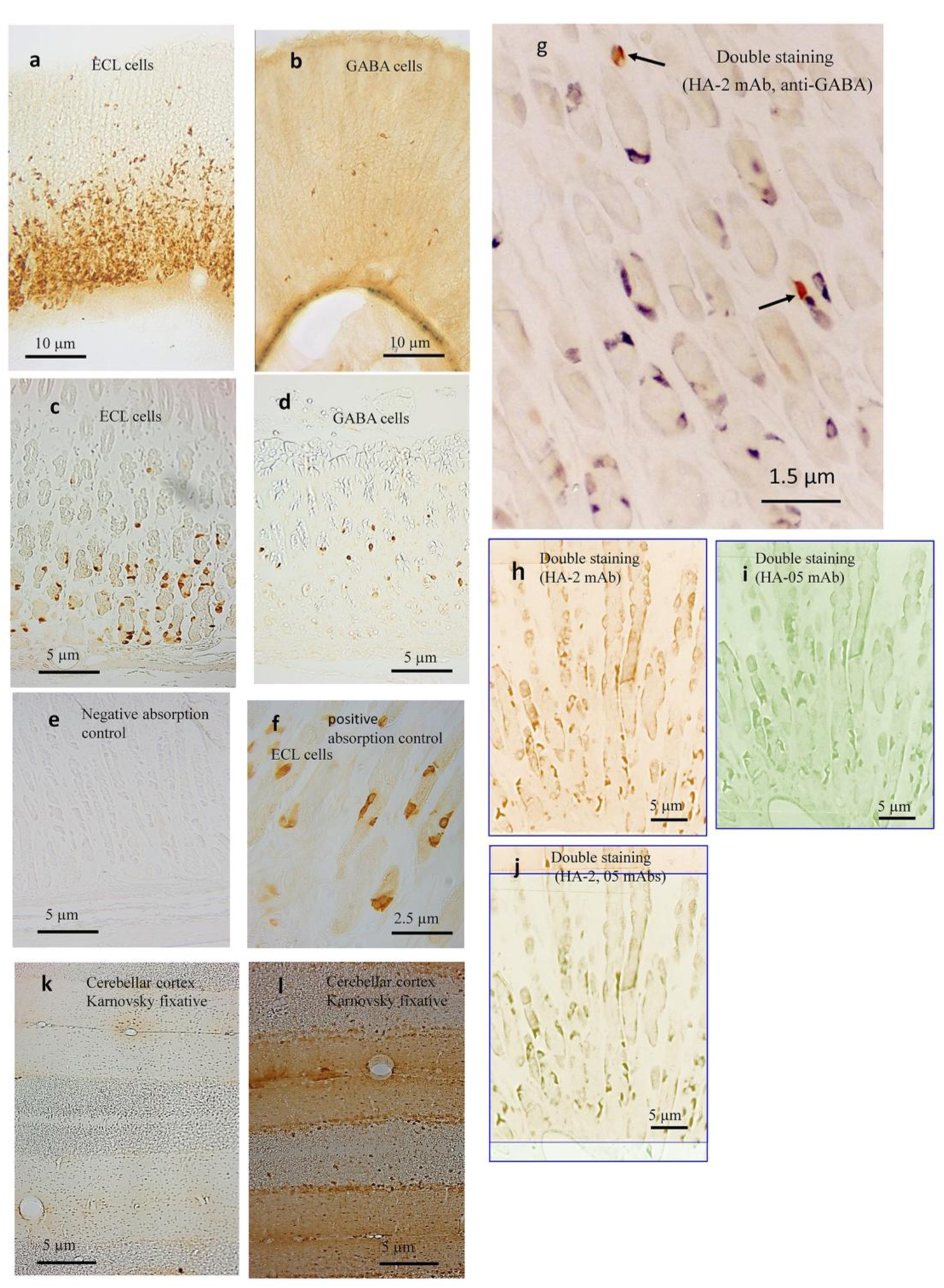
Verification of HA2 antibody specificity and fixative dependency. Because histamine and GABA exhibited similar staining patterns in the brain, antibody specificity was verified in gastric fundus tissue, where the two cell populations are clearly distinct. Absorption controls, dual labeling, independent antibody validation, and fixative dependency confirmed the specificity of HA2. **(a)** HA2 selectively labels enterochromaffin-like (ECL) cells clustered in the lower gastric glands. **(b)** GABA mAb labels a sparse, morphologically distinct cell population with thin apical processes. **(c, d)** High-magnification paraffin sections demonstrate the distinct morphologies of histamine-positive ECL cells **(c)** and GABA-positive cells **(d)**. **(e, f)** Preabsorption with histamine–GA–BSA abolishes HA2 immunoreactivity (*6–9,18*), whereas GABA–GA–BSA has no effect. **(g)** HA2 and GABA mAbs label non-overlapping cell populations. **(h–j)** Independent validation using HA2 (**h**) and HA05 (**i**) demonstrates identical histamine labeling patterns (*6,18*). **(k, l)** Karnovsky fixation (*34*) abolishes HA2 labeling by altering the GA-dependent histamine epitope, whereas GABA immunoreactivity is preserved. **Scale bars:** 10 μm **(a, b);** 5 μm **(c–e, h–j, k, l);** 2.5 μm **(f);** 1.5 μm **(g).**

**Extended data Figure 2.**
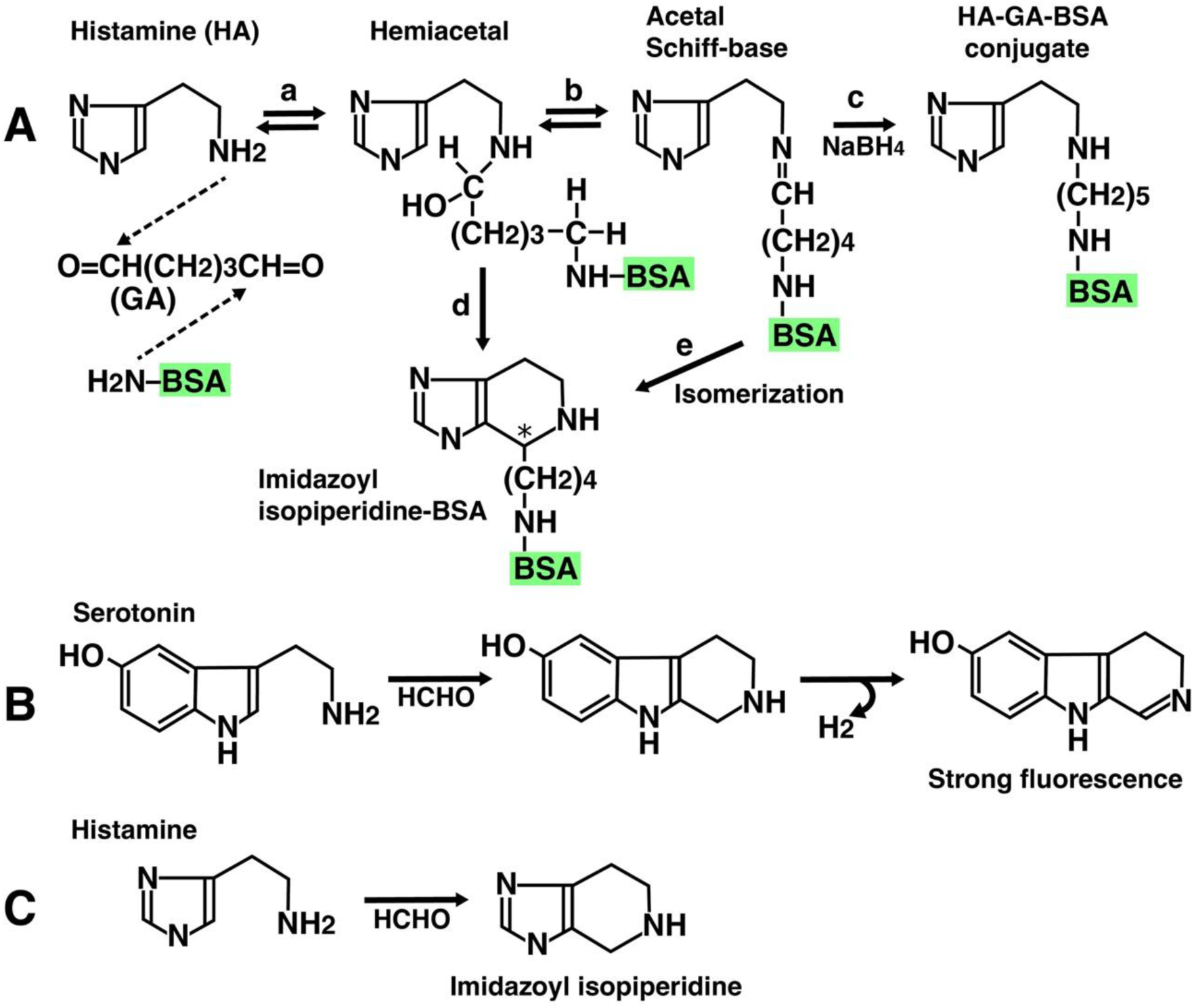
Presumed chemical structure of glutaraldehyde-dependent histamine epitope. All five monoclonal antibodies (HA1-HA5) recognize histamine in gastric ECL cells (*6–9*), but only HA2 detects histamine in the brain. This difference may be explained by our previous study that the histamine-GA-BSA antigen has at least two different epitope structures (*18*). The chemical structures of these hypotheses are presented below. (**A**) Histamine reacts with the aldehyde group at one end of glutaraldehyde to form a hemiacetal, and the reaction proceeds in two directions. In one pathway, a Schiff base is formed, and a histamine-protein complex stabilized with NaBH4, as described in Methods. In the other pathway, the hemiacetal and histamine itself condense to produce an imidazolyl isopiperidine derivative. Furthermore, the Schiff base isomerizes to imidazolyl isopiperidine. This cyclic structure may preferentially form under intracellular conditions within nerve cells, as demonstrated by Falck-Hillarp reaction (*35*). (**B**) Serotonin undergoes a Falck-Hillarp-type aldehyde condensation reaction, producing a cyclic piperidine fluorescent product (*35*). (**C**) Histamine similarly reacts with formaldehyde to produce an imidazolyl isopiperidine structure similar to the classical monoamine-aldehyde reaction (*36*).

## Methods and Materials

### Animals and Ethical Statement

Adult male Wistar rats (200–300 g; Kyudo Experimental Animals, Kumamoto, Japan) were housed at 21 ± 1 °C under a 12-h light/dark cycle with food and water available *ad libitum*. All procedures complied with national guidelines and were approved by the Sojo University Ethics Review Committee for Animal Experimentation.

### Tissue Preparation and Fixation

Rats were deeply anesthetized with sodium pentobarbital (60 mg/kg, i.p.) and transcardially perfused with phosphate-buffered saline (50 ml/min, 2 min), followed by freshly prepared 5% glutaraldehyde (GA) in 10 mM phosphate buffer (pH 7.2) for 6 min. Brains were removed, post-fixed overnight at 4 °C in the same fixative, and sectioned at 50 μm using a Microslicer (Dosaka EM, Kyoto, Japan). Alternate sections were processed for histamine (HA) or GABA immunohistochemistry.

### Immunohistochemistry (IHC)

IHC was performed as previously described (6–9,*15–18*). To eliminate non-specific background from residual reactive aldehydes, all sections were treated with 0.2% NaBH₄ for 10 min.

For histamine (HA) staining, free-floating sections were incubated for 48 h at 4 °C with the monoclonal antibody HA2 (1:500–1:1,000) in 50 mM Tris-HCl-buffered saline (TBS, pH 7.4) containing 0.1% saponin, 0.25% bovine serum albumin (BSA), and 1% normal goat serum. Sodium metabisulfite (0.1% Na₂S₂O₅) was included to stabilize the antigen. Sections were then incubated with HRP-conjugated goat anti-mouse IgG/Fab′ (1:200; MBL, Nagoya, Japan) for 12 h at 4 °C, and immunoreactivity was visualized with diaminobenzidine (DAB) and H₂O₂.

GABA immunostaining was performed using the same protocol with a monoclonal anti-GABA antibody (Funakoshi, Japan). For 5-μm paraffin sections of gastric tissue, protease digestion (Type XXIV bacterial protease) was included to enhance antibody penetration (6–9).

### Immunoelectron Microscopy

Following the free-floating DAB reaction, specimens were post-fixed with 1.0% osmium tetroxide in 50 mM cacodylate buffer (pH 7.4) for 1 h. Tissues were dehydrated through a graded ethanol series, cleared in propylene oxide, and embedded in Epon 812 resin. Regions of interest were excised using a 2-mm diameter punch, mounted on Epon blocks, and processed into ultrathin sections. Sections were carbon-coated and examined using a 100CX electron microscope (JEOL, Tokyo, Japan) (*7–9, 15, 17*).

### EM Quantification of Vesicular Core Localization

Tissue samples were processed for pre-embedding DAB immunoperoxidase labeling (HA2), post-fixed with osmium tetroxide, embedded in Epon, and cut into ultrathin sections (60–70 nm) for transmission electron microscopy. Only intact, non-overlapping vesicles were analyzed, with immunonegative asymmetric synapses serving as internal controls. DAB intensity was measured along radial transects and normalized to the vesicle radius (0–100%). The vesicle core was defined as 0–50% of the radius, and the peripheral region as 50–100%. Across 23 boutons, a total of 2,195 core and 108 peripheral DAB signals were quantified, yielding a core fraction of 95.1%. Statistical significance was assessed by permutation analysis (10,000 iterations) using randomized radial distributions as the null model, and bootstrap resampling (10,000 iterations) was performed to estimate confidence intervals.

### Double-Staining Immunohistochemistry

Co-localization was examined by sequential immunostaining as previously described (*29*). Histamine was first visualized with 4-chloro-1-naphthol (0.02% in TBS containing 0.005% H₂O₂ and 2% ethanol). After digital imaging, bound antibodies were stripped with 0.1 M glycine-HCl (pH 2.2; four 30-min incubations), followed by ethanol decolorization. Sections were then blocked overnight and reprocessed for GABA immunohistochemistry using the DAB method.

### Specificity and Control Experiments

Specificity was verified by (i) omission of the primary antibody, (ii) incubation with an isotype-matched irrelevant monoclonal antibody (anti-penicillin IgG1, 30–80 ng/ml), and (iii) preabsorption of HA2 with histamine–GA–BSA conjugate (5 μg/ml). All controls abolished or excluded specific HA2 immunoreactivity, confirming its high specificity for GA-fixed histamine.

### Single-nucleus RNA sequencing (snRNA-seq) Analysis of Cerebellar Subclasses

To investigate the expression profile of *Hdc* across distinct cerebellar cell populations, a single-nucleus RNA sequencing (snRNA-seq) dataset generated by the Macosko laboratory (*37*) was analyzed. The dataset, obtained through the Allen Brain Cell (ABC) Atlas, contains transcriptomic profiles of adult mouse brain nuclei, classified according to the mouse brain taxonomy.

Downstream bioinformatic analysis was performed using Scanpy (v1.11.5). For cell type identification, the high-level annotation metadata provided by the original repository was utilized, focusing on four primary cerebellar subclasses: Granule cells, Molecular Layer Interneurons expressing Cdh22 (MLI:Cdh22), Molecular Layer Interneurons expressing Megf11 (MLI:Megf11), and Purkinje cells.

Gene expression distributions across individual cells in each subclass were visualized using violin plots. The normalized expression values [log2(CPM + 1)] were used to plot the density and distribution for each cell type. To calculate the proportion of *Hdc*-positive [*Hdc*(+)] cells within each subclass, cells with an expression value strictly greater than zero for the *Hdc* transcript were defined as *Hdc*(+) cells. The percentage of these positive cells relative to the total number of cells within each respective subclass was calculated and visualized using a bar chart.

## Ethics statement

Ethics approval for this study was obtained from the Sojo University Animal Experiment Ethics Committee.

## Authors contributions

F.K. designed the experiment. F.K. performed the experiments. F. K. analyzed the data. F. K. drew the photographs and figures. F.K. wrote the manuscript.

## Abbreviations

IHC: Immunohistochemistry
GABA: γ-aminobutyric acid
NaBH₄: sodium borohydride
HDC: histidine decarboxylase
HA2: glutaraldehyde-dependent anti-histamine monoclonal antibody (mAb)
HA05: anti-histamine monoclonal antibody (mAb)
GA: glutaraldehyde
TMN: tuberomammillary nucleus
CNS: central nervous system
ECL: enterochromaffin-like cell
EM: electron microscopy
LM: light microscopy
DAB: 3,3′-diaminobenzidine
IP: immunopositive
IN: immunonegative
VGAT: vesicular GABA transporter
VMAT: vesicular monoamine transporter
*Hdc*: histidine decarboxylase transcripts

## Acknowledgements

I would like to express my deep gratitude to my wife, the late Chiemi Fujiwara, who supported me both materially and emotionally. I would also like to express my deep gratitude to the late Takashi Suemasu, who helped me take the electron microscope photographs, to Professor G. Bay, K. Maeda, M. Matsumoto, and C. Tamura, who helped me with my research, to Professor S. Matsufuji for valuable discussion, to Dr. Yu Liu, who helped me with the Allen analysis, to Y. Shirahama, who helped me with my computer, and to H. Fujiwara, M. Tateishi, and K. Tateishi, who provided their kind support. I am truly grateful.

## Competing interests

The author declares no competing interests.

**Correspondence** should be addressed to Kunio Fujiwara

